# Mites alight! Sunflower crop area and pollen supplementation enhance honey bee resistance to *Varroa destructor*

**DOI:** 10.1101/2021.08.04.454914

**Authors:** Evan C Palmer-Young, Rosemary Malfi, Yujun Zhou, Bryanna Joyce, Hannah Whitehead, Jennifer I Van Wyk, Kathy Baylis, Kyle Grubbs, Dawn Lopez, Jay D Evans, Rebecca E Irwin, Lynn S Adler

## Abstract

Anthropogenic landscape changes can affect parasite epidemiology in wild and agricultural animals. Honey bees are agricultural animals whose services are threatened by loss of floral resources and by parasites, most notably the invasive mite *Varroa destructor*. Existing mite control strategies rely heavily on chemical treatments that can adversely affect bees. Alternative, pesticide-free control methods are urgently needed to maintain effective pollination services. Many flowering plants provide nectar and pollen that enhance resistance to parasites in animals. Enrichment of landscapes with antiparasitic floral resources could therefore provide a sustainable means of parasite control in pollinators.

Floral rewards of Asteraceae plants have been shown to reduce parasitic infection in diverse bee species, including honey and bumble bees. Here, we tested the effects of sunflower (*Helianthus annuus*) cropland and pollen supplementation on honey bee resistance to macro- and microparasites. Our results show that each doubling of sunflower crop area is associated with a 28% reduction in mite infestation intensity. Late-summer supplementation of colonies with sunflower pollen reduced mite infestation by 2.75-fold relative to an artificial pollen. Our findings suggest the potential for sunflower plantings or pollen supplements to counteract a main driver of honey bee losses worldwide.

## INTRODUCTION

Nutritional resources have profound effects on host-parasite interactions [1]. The nutrition of free-ranging animals is shaped by foraging opportunities in local landscapes, which vary in resource quality, quantity, and medicinal potential. As landscapes become increasingly human-dominated and agriculturally intensified, foraging patterns of animals—including insects in agroecosystems—are heavily influenced by anthropogenic land use change [2]. Plant-pollinator communities appear particularly vulnerable to both parasites—which spread easily given the broad foraging ranges, high activity levels, and human-assisted migrations of social bees [3]—and to land use changes, which govern the floral food resources and habitat upon which survival and reproduction depend [4].

Due to their worldwide distribution, high densities, and intimate association with agriculture, honey bees provide an exemplary system for investigating how land use-mediated changes in floral resources affect host-parasite dynamics. The free-ranging, semi-domesticated European honey bee (*Apis mellifera*) depends on floral resources—including those in cropland—for nectar and pollen. In return, managed honey bees provide roughly half of all visits to flowers and the majority of pollination in cropland [5]—a service that increases yields in the majority of crops [6] and is valued at roughly 10% of global agricultural production [7]. These pollination services are threatened by land use-driven reductions in availability of floral resources [4,8,9], which can compromise the nutrition and physiology of individuals and colonies [10–12]. Honey bees are also infected by a suite of parasites and pathogens that contribute to colony loss and pose threats of spillover into wild bee communities [13,14]. Parasites implicated in colony mortality include the microsporidian *Nosema ceranae*, viruses, and the virus-vectoring ectoparasitic mite *Varroa destructor* [13,15].

Resistance and tolerance of bees to parasites is shaped by nutrition derived from floral nectar and pollen. While nectar and the resulting honey provide a sugar-rich source of energy, dietary pollen provides amino acids and lipids that support overall development, tolerance to parasites, and immune system activity [16–18]. Beyond supplying macronutrients, specific floral and other plant resources can enhance resistance to infections [19]. Plant resin-derived propolis has a variety of immunomodulatory and antimicrobial effects, protecting colonies from the abiotic environment, predation, and parasitic infection [20]. Nectar and pollen also supply a diverse assortment of plant secondary metabolites, including many compounds with effects on insect parasites [21], immunity [22], and infection [23,24]. At a broader scale, the structure of floral landscapes and the presence of specific species that affect parasite transmission can affect the prevalence of infection in bee communities [25–28]. Loss of antiparasitic plant resources could exacerbate susceptibility to infection, whereas increased plantings could enhance disease resistance in wild and managed bees [24].

One plant species of particular interest for bee health is sunflower (*Helianthus annuus*, Asteraceae). Its abundant production of nectar and pollen is critical for the reproduction of numerous wild bee species in the plant’s native North American range [29,30]. Sunflowers are also cultivated plants that constitute major oilseed crops in Europe and Asia—which combine for 68% of global production [31]—as well as the Americas, making important contributions to late-season pollen and honey storage in honey bee colonies co-located in agricultural landscapes [32,33]. However, sunflower is generally not considered a preferred resource for honey bees [30,34,35]. Like that of many Asteraceae, the pollen of sunflower is low in protein compared to other bee-visited plants, and exclusively sunflower-based diets do not support development of honey and bumble bees [35–37]. This has led to the conclusion that Asteraceae pollen is an inappropriate food source for bees that lack special physiological adaptations to its secondary metabolites and nutritional deficiencies [37].

On the other hand, several studies have demonstrated benefits of nectar and pollen from sunflower and other Asteraceae for resistance to parasites. Asteraceae pollen provisions were associated with resistance to brood parasites in *Osmia* [38]. Pollen of sunflower and *Solidago* spp. (Asteraceae) afforded resistance to gut infection with the trypanosomatid parasite *Crithidia bombi* in the bumble bee *Bombus impatiens* [39], with sunflower cropland associated with decreased infection intensity at the landscape scale [40]. In honey bees, consumption of sunflower pollen and honey reduced infection by the gut microsporidian *Nosema cerana*e [40]—a parasite implicated in colony collapse [41]—with nurse bees exhibiting infection-induced preference for sunflower honey [42].

These findings with individual bees led us to hypothesize that the availability of sunflower resources could also reduce levels of parasites or pathogens in honey bees at the colony and landscape scale. To test this hypothesis, we combined a multi-year, nationwide landscape survey with field and lab experiments to evaluate the effects of sunflower cropland and pollen diets on resistance of honey bees to micro- and macroparasites associated with colony loss [13,15], including viruses, gut microsporidia in the genus *Nosema*, and the ectoparasitic mite *Varroa destructor* (hereafter *“Varroa*”). We had particular interest in *Varroa* for three reasons: first, because of its central role in colony loss and vectoring of other pathogens [15,43]; second, due to the coincidence of seasonal late-summer increases in infestation [44] with the timing of sunflower bloom [45]; and third, due to its biology as a brood-dependent macroparasite, against which Asteraceae pollen diets provided protection in other bee hosts [38].

We first used data from the US Department of Agriculture’s Animal and Plant Health Inspection Service (USDA-APHIS) Honey Bee Disease Survey [46] to test for associations between sunflower crop area and *Nosema* and *Varroa* in co-located colonies. We then conducted field experiments in spring and late summer to evaluate the effects of sunflower pollen-based supplements on *Nosema, Varroa*, and viruses in colonies. In addition, we used a lab experiment with caged worker bees to evaluate the effects of a sunflower pollen-based diet on naturally occurring viral infection under controlled conditions. Lastly, to illustrate temporal trends in honey bee access to this putatively antiparasitic floral resource, we quantified changes in US cropland planted with sunflowers over the past 40 years. Our study provides the first evidence for crop composition-mediated change in *Varroa* resistance in honey bee colonies, suggesting a tangible strategy for reducing infestation-mediated colony losses in a preeminent agricultural pollinator.

## MATERIALS AND METHODS

We briefly summarize each experiment below. For full details, please refer to Supplementary Information: Extended Methods.

### Landscape associations between sunflower, *Varroa*, and *Nosema*

We used data from the USDA-APHIS National Honey Bee Disease Survey—which assesses *Varroa* and *Nosema* levels in colonies of >400 apiaries in >30 US states each year [46]—to evaluate associations between sunflower crop area and bee infections. Sunflower crop area within a 2-mile (3.2 km) radius of the apiary was quantified using the National Agricultural Statistics Service (NASS) cropland data layer (retrieved from nass.usda.gov) with 30 m^2^ pixel resolution. Land cover within this radius (8000 acres) encompasses the typical foraging range of honey bees [47] and has been shown to correlate with colony health metrics [48]. We obtained meteorological covariates (average minimum temperature (°C) and precipitation (mm) during the collection month within 3.2 km of the apiary) from Oregon State University’s Prism database (https://prism.oregonstate.edu/).

The original APHIS dataset contained 2687 observations of colonies between 2010 and 2015. We restricted our analyses to observations made in August and September, the period during and immediately following sunflower bloom in temperate climates [45]. Observations were filtered to remove states for which there were < 3 sites with any sunflower cropland within 3.2 km of the apiary. This retained 181 observations from 5 states, of which 48 (26.5%) were associated with sunflower: SD (n = 20 of 57), ND (n = 12 of 19), NY (n = 7 of 28), GA (n = 6 of 49), and WI (n = 3 of 28), with a median non-zero sunflower coverage of 25.5 acres (i.e., 0.32% of total land cover, interquartile range: 0.015 to 3.1%).

Associations between sunflower crop area and *Varroa* mite infestation (measured as mites per 100 bees, using subsamples from 8 colonies per apiary [46]) and *Nosema* infection intensity (in 10^6^ spores per bee) were tested using negative binomial family linear mixed models [49] with log-transformed sunflower acreage as a continuous predictor and US state as a random effect. The *Varroa* model included average minimum monthly temperature and the *Nosema* model included month of collection as additional predictors. We excluded month of collection from the *Varroa* model (Z = 0.94, P = 0.35) and average minimum temperature from the *Nosema* model (Z = −1.61, P = 0.10), as these terms did not explain significant variation in the response variables. We initially attempted to include year of collection and average precipitation as additional predictors; however, the addition of these variables resulted in convergence warnings (indicative of model overfitting), so they were excluded from the final model. This and all subsequent analyses were performed using R v4.0 for Windows [50].

### Effects of late-summer sunflower pollen supplementation on field colonies in Maryland

#### Experimental design

A cohort of 30 colonies—all in the same apiary (College Park, MD USA)— was supplemented weekly for five weeks (4 September through 9 October 2019) with a 900 g patty of sunflower, wildflower, or artificial pollen (“BeePro”, Mann Lake Beekeeping, Wilkes-Barre, PA USA; n = 10 colonies per treatment). We assessed *Varroa* mite infestation and infection with *Nosema* and seven viruses (Acute Bee Paralysis Virus (ABPV), Chronic Bee Paralysis Virus (CBPV), Deformed Wing Virus (DWV), Israeli Acute Paralysis Virus (IAPV), Kashmir Bee Virus (KBV), Lake Sinai Virus-2 (LSBV-2), and Varroa destructor Virus-1 (VDV-1)) on the first and last days of the supplementation period using the methods of the National Honey Bee Disease Survey [46].

#### Statistical analyses

Differences in consumption of the three pollen treatments across treatments and over the five weekly collections were tested with a linear model using proportional consumption of the patty (as estimated by mass loss) as the response variable and pollen treatment, time, and the treatment by time interaction as predictor variables. *Varroa* infestation was analyzed with a Poisson family generalized linear model [50] with pollen treatment as a fixed effect. We excluded three colonies (one in each treatment) with high initial infestation (>2 mites per 100 bees, more than twice as high as in any of the 27 other colonies). Because infestation intensity (<1 mite per 100 bees) and variance in the remaining colonies were negligible at the start of the experiment, only the endpoint infestation levels were analyzed. Intensities of infection with DWV and VDV-1 were analyzed using a negative binomial model with treatment, time point (i.e., pre- and post-treatment) and their interaction as predictor variables and a random intercept for each colony to account for repeated measures. We used the treatment by time interaction term to evaluate whether changes in infection intensity over the course of the experiment differed across pollen treatments. The other viruses were detected rarely (IAPV, 8.3% and CBPV, 1.7%) or not at all (ABPV, KBV, LSV-2), precluding statistical analysis. *Nosema* infection intensity was low, with only 2 of 60 samples exceeding the typical treatment threshold of 10^6^ spores per bee. Prevalence (detectable spores in only 30% of colonies) was too low for convergence of binomial mixed models given the sample size, so this parasite was also not analyzed.

### Effects of springtime sunflower pollen supplementation on field colonies in Massachusetts

Field colonies owned and managed by a commercial beekeeper (N = 11 per treatment, divided among five sites) were supplemented for five weeks (23 May-27 June 2018) with a 170 g patty containing one of four organic bee pollen treatments: sunflower (approximately 88% sunflower and 12% wildflower), wildflower (approximately 11% sunflower and 89% wildflower), sunflower-wildflower mix (approximately 28% sunflower and 72% wildflower), or artificial pollen (BeePro). A fifth treatment group received no supplementation. *Varroa* and *Nosema* were measured pre- and post-treatment (0 and 5 weeks) by alcohol washes and microscopic spore counts, respectively. Viruses were assessed post-treatment only, using the same methods and targets as in the late-summer experiment [46].

#### Statistical analysis

*Varroa* mites were detectable in only one-third of samples, with trivial infestation (median 0, mean 0.19 mites per 100 bees), and were not further analyzed. *Nosema*—though found at 60-90% prevalence at baseline—was likewise rare at the post-treatment follow-up (median spore count of zero in all treatments), and not analyzed.

We analyzed infection intensity for the three viruses with >50% prevalence—VDV-1 (100% of 43 colonies), DWV (77%), and LSV-22 (53%). The four viruses detected at <20% prevalence (CBPV and KBV: each found in 7 of 43 colonies (16%); IAPV: 2 of 43 colonies (4.7%); ABPV: not detected) were not further analyzed. Negative binomial models with site as a random effect did not converge. We therefore fit separate mixed-effects models [51] for prevalence (binomial model on presence-absence data) and infection intensity (log-normal model on samples with measurable infection). For VDV (100% prevalence), the binomial model was omitted.

### Effects of sunflower pollen on caged bees

Individual plastic cups [52] were established with 30 mature worker bees each, collected from a single colony in the Beltsville, MD Bee Research Laboratory research apiary (*Apis mellifera ligustica)* in April 2018. This colony had endemic infections of DWV and *Nosema ceranae*. Six cups were established under each of five treatments—a pollen-free control group plus 4 pollen treatments containing sunflower and wildflower pollen (Koppert Biological Systems, Howell, MI, USA) in ratios of 3:0, 2:1, 1:2, and 0:3. Cups were maintained for six days in a dark incubator at 34°C. Bulk RNA extractions of 20 bees per cup were used to quantify infection with DWV, VDV-1, *Nosema ceranae*, and trypanosomatids. Infection intensities were normalized to mRNA levels of the honey bee reference genes actin and RPS5 [52].

Normalized infection intensities were estimated using ΔCt (i.e., log_2_ scale) by subtracting the cycle time (Ct) values of the target from the mean Ct of the two normalizer genes (actin and RpS5) for the corresponding sample [52]. Target samples that did not pass the fluorescence threshold were arbitrarily given a Ct of 40. Samples for which Ct values differed by >5 cycles between the two technical replicates were excluded from analysis, as were two samples for which the Ct values of actin and RpS5 were strongly discordant. This removed two samples each from the all-wildflower pollen treatment (both DWV and VDV-1) and the 67% and 100% sunflower treatments (DWV only). Differences in infection intensity (i.e., ΔCt score) across treatments were tested using analyses of variance in R [50]. *Nosema ceranae* was detected in only one sample and trypanosomatids in only 4 samples (all in trace quantities with Ct >34.6); these responses were not further analyzed.

### Longitudinal changes in US sunflower crop area

To quantify long-term trends in honey bee access to sunflower cropland, NASS crop records were used to assess changes in sunflower crop area nationwide over the last 40 years. Total area planted was modeled as a function of time using a generalized linear model with Poisson error distribution. Based on visually apparent trends, separate models were fit to the entire time series (1980-2021) and the most recent data (1995-2021).

## RESULTS

### Landscape associations between sunflower, *Varroa*, and *Nosema*

*Varroa* mite infestation was negatively associated with sunflower crop area (χ^2^_1_ = 18.86, P < 0.001, Fig. 1) and average minimum temperature (χ^2^_1_ = 13.38, P < 0.001). Models predicted a 28% decrease in mite infestation (coefficient for log-transformed area: −0.28 ± 0.065 SE, Z = −4.34, P < 0.001) for each doubling of sunflower crop area and a 6% increase for every 1 °C decrease in minimum temperature (coefficient: −0.062 ± 0.017 SE, Z = −3.66, P < 0.001). This is consistent with typical rises in infestation as the weather cools during the late summer and early fall. *Nosema* infection intensity was 63% lower in September than in August (coefficient: −0.63 ± 0.24 SE, Z = −2.61, P = 0.009), but not correlated with sunflower crop area (Z = −0.63, P = 0.53).

**Figure 1.**
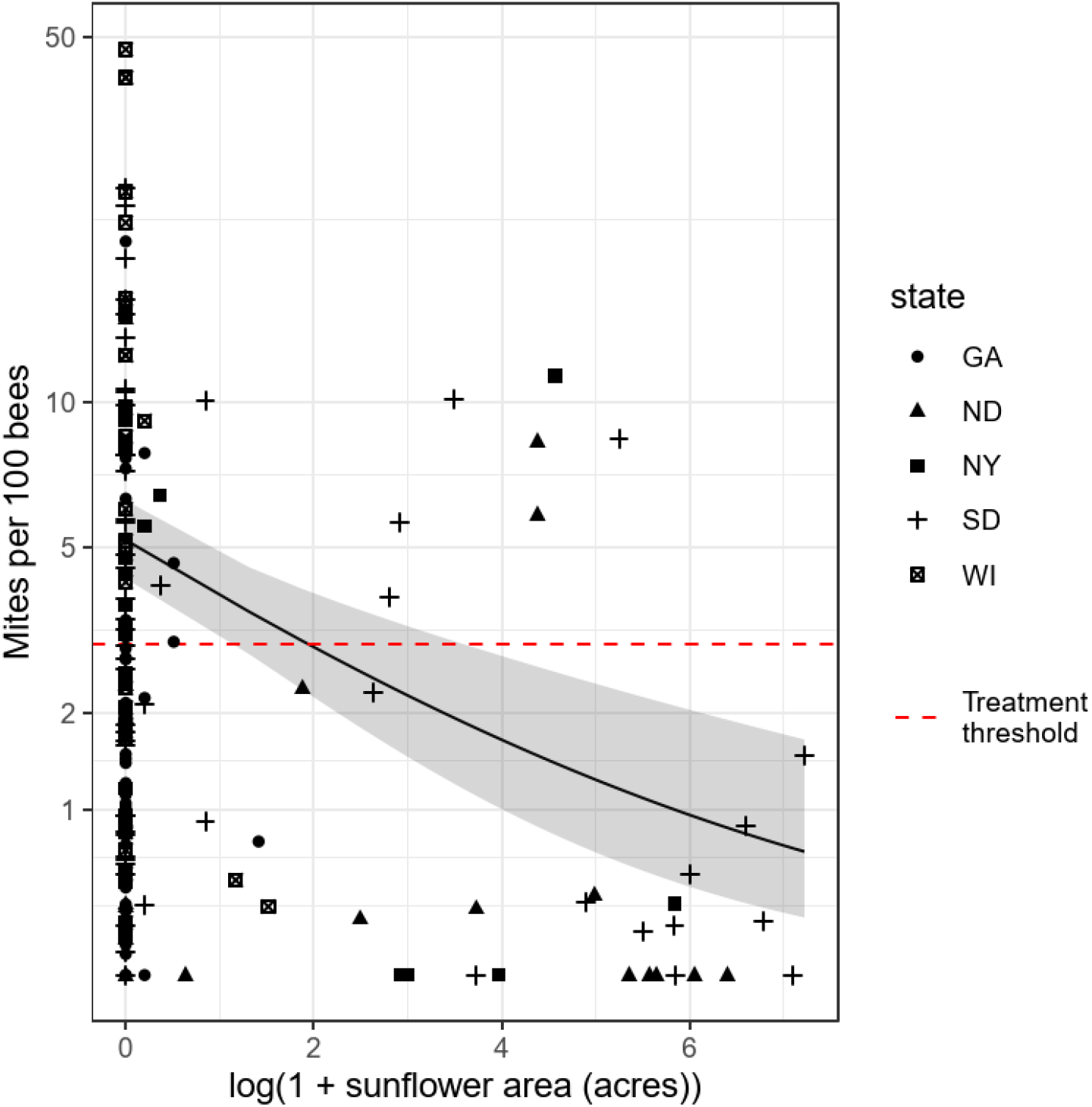
Association between sunflower crop area and *Varroa* mite infestation. X-axis shows natural log-transformed sunflower crop area within a 3.2 km radius. Y-axis shows infestation intensity in mites per 100 bees. Points represent samples from different apiaries. Trendline and shaded band show model predictions and standard errors from the fixed effects portion of the fitted model at an average value of the covariate (average minimum temperature). Dashed line indicates canonical treatment threshold (3 mites per 100 bees). Data are from the USDA APHIS National Honey Bee Disease Survey 2010-2015, filtered to include only data in August and September (the months immediately following sunflower bloom) from states with at least 3 apiaries associated with sunflower crops during the chosen time window. Shapes indicate US states in which sampled apiaries were located (GA: Georgia, ND: North Dakota, NY: New York, SD: South Dakota, WI: Wisconsin).

### Effects of late-summer sunflower pollen supplementation on field colonies in Maryland

#### Pollen consumption

Pollen treatments were consumed by the experimental colonies, with an average consumption of 686 g (76% of the 900 g patty) per week. Mean proportions consumed differed across pollen treatments (F_2,114_ = 6.60, P = 0.002), but by <20% (Sunflower: 70%, BeePro: 77%, Wildflower: 82%). Proportions consumed declined over the 5 weeks of sampling (F_1,114_ = 22.9, P < 0.001), from an average of 87% during the first week to 67% during the final week. However, this decline did not vary significantly by pollen treatment (treatment x time interaction: F_2,114_ = 0.73, P = 0.48).

#### *Varroa* infestation

Post-supplementation *Varroa* infestation levels differed across the pollen treatments (χ^2^_2_ = 11.83, P = 0.003, Fig. 2). There was a 2.75-fold reduction in infestation intensity in the sunflower vs. the BeePro artificial pollen treatment (Z = 3.23, Tukey-adjusted P = 0.004). Infestation in the wildflower treatment (3.02 ± 0.58 SE mites per 100 bees) was intermediate between that of the sunflower (1.54 ± 0.41 SE) and the BeePro treatment (4.23 ± 4.23 SE), but not significantly different from either the sunflower treatment (Z = 2.04, Tukey-adjusted P = 0.10) or the BeePro treatment (Z = 1.35, Tukey-adjusted P = 0.37).

**Figure 2.**
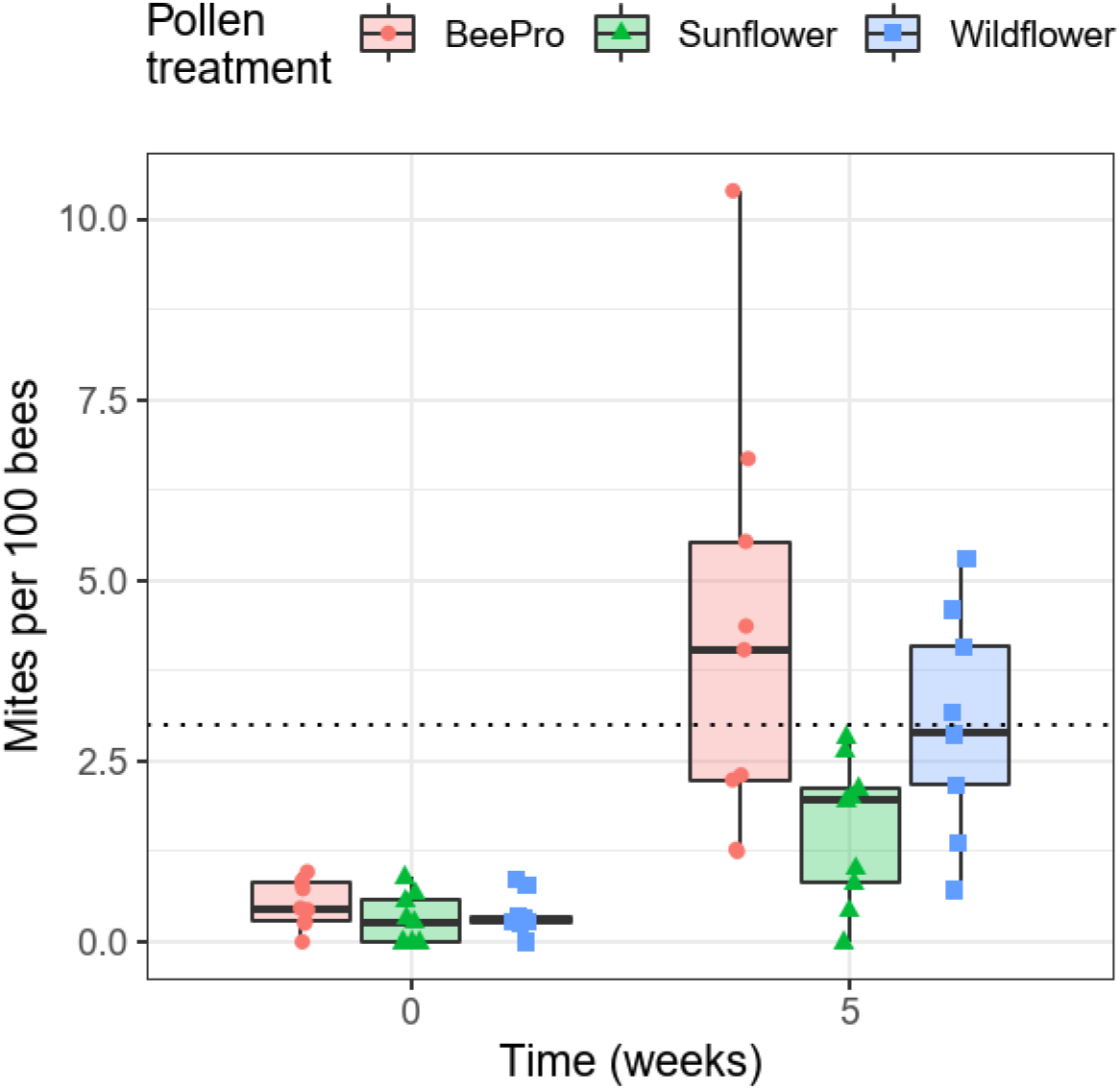
Effects of sunflower pollen supplementation on *Varroa* mite infestation of field colonies. X-axis shows collection dates (before and after 5 weeks of supplementation). Y-axis shows infestation intensity in mites per 100 bees. Points represent samples from different colonies. Boxplots show medians and interquartile ranges. Dashed line indicates canonical treatment threshold (3 mites per 100 bees).

#### Viruses

VDV-1 was the most prevalent virus surveyed, detected in all but one of the 60 samples. Infection intensity did not vary across treatments (χ^2^_2_ = 1.87 P = 0.39), collection dates (χ^2^_1_ = 1.17, P = 0.28), or their interaction (χ^2^_2_ = 0.17, P = 0.92). DWV was the second most prevalent virus, found in 88% of samples. Only collection date was a significant predictor of infection (χ^2^_1_ = 5.11, P = 0.024), with 90% higher infection intensity at the end vs. the start of the experiment (Z = 2.22, P = 0.03). Infection by DWV did not differ significantly by pollen treatment (χ^2^_2_ = 0.21, P = 0.89), nor was there a significant treatment by time interaction (χ^2^_2_ = 1.15, P = 0.56).

### Effects of springtime sunflower pollen supplementation on field colonies in Massachusetts

All pollen treatments were consumed by the experimental colonies (mean weekly consumption ± SD: 93% ± 16%), although the artificial pollen (BeePro, mean 73%) appeared somewhat less palatable than the honey bee pollens (pooled mean 98%). No significant effects of pollen treatment were found for virus prevalence (DWV: χ^2^_3_ = 2.46, P = 0.48; LSV-2: χ^2^_3_ = 0.46, P = 0.93) or intensity (VDV-1: χ^2^_3_ = 0.74, P = 0.86; DWV: χ^2^_3_ = 5.17, P = 0.16; LSV-2: χ^2^_3_ = 5.06, P = 0.17).

### Effects of sunflower pollen on caged bees

No significant effects of pollen treatment were found for infection with DWV (F_4, 19_ = 0.49, P = 0.74) or VDV-1 (F_4, 23_ = 0.67, P = 0.62).

### Longitudinal changes in US sunflower crop area

NASS records showed that sunflower crop acreage planted in the USA has declined by an average of 2.03% per year since 1980 (Poisson model estimate: −2.03 · 10^−2^ ± 8.3 · 10^−6^ SE, Z = −2446, P < 0.001). An even steeper decline (−3.55% per year) was found since 1995 (estimate: −3.55 · 10^−2^ ± 1.71 · 10^−5^ SE, Z = −2073, P < 0.001).

## DISCUSSION

We found negative associations between sunflower cropland and *Varroa* infestation, suggesting that floral resources and agricultural decisions can shape not only bee nutritional status, but also resistance to one of the most formidable parasites of managed honey bees worldwide. Our experimental manipulations of field colonies indicate a causal role for sunflower in reducing infestation, whereas our analyses of crop records suggest that honey bee access to sunflower cropland is declining.

Originally a parasite of *Apis cerana, Varroa destructor* has become ubiquitous in *Apis mellifera* colonies on all continents except Australia [44,53]. Infestation intensity has been consistently implicated in colony mortality [53], depleting the nutritional reserves of pupae and adult bees and exacerbating the transmission and virulence of otherwise asymptomatic viral infections [43,54], including those implicated in wild bee mortality [14]. Infestation of honey bees therefore presents concerns for crop pollination services and bee conservation alike.

Current *Varroa* treatments are largely based on chemical acaracides—typically applied three times per year [12]—that can suppress honey bee immune function [55], render contaminated hive products unsuitable for human consumption [56], and lose efficacy as mites evolve resistance [53]. Pollen consumption can improve tolerance of honey bees to infestation [16,17]. However, studies that show effects of land use and floral resource availability on colony health have yet to implicate specific plant species in resistance to infestation [8,12]. Instead, the overwhelming effects of infestation on colony survival have been considered to obscure—rather than highlight—the benefits of resource-rich landscapes [8]. Our results draw a connection between the availability of a specific floral resource and infestation intensity, with models predicting a 28% decrease in infestation intensity for every doubling of sunflower crop area (Fig. 1). This finding adds to the body of research on antiparasitic effects of sunflower nectar and pollen in other bee hosts and against other infections in honey bees [39,40,42] and builds on previous findings of Asteraceae pollen-mediated resistance to brood parasites in solitary bees [38]. Our results are also consistent with an association between use of colonies for sunflower pollination and low *Varroa* infestation intensities in India [57].

Our experiments with field colonies indicate that sunflower pollen can cause reductions in *Varroa* infestation. The 5-week time period over which this effect was achieved corresponds to both the period immediately following sunflower bloom [45] and the time when *Varroa* infestations begin to rise towards seasonal peaks in late autumn [44]. This effect was discernible (Fig. 2) despite the cohabitation of all treated colonies in the same apiary, which could have obscured treatment effects by allowing drift of bees and their mites between colonies [56], and while allowing bees to forage freely on other pollen sources. Although application of pollen traps or removal of pollen stores would have accentuated differences between diets across treatments, our intention was to simulate the availability of sunflower within the context of a landscape and growing season. Given that honey bees in many areas do not normally visit sunflowers [30,34], but may exhibit infection-induced changes in foraging [59]—including *Nosema-*induced preference for sunflower honey [42] and *Varroa-*related collection of propolis [60]— infestation-induced preference for sunflower seem plausible. However, expression of such behavior would depend on sunflower forage availability.

The mechanisms underlying the effects of sunflower on *Varroa* infestation remain unresolved, but could reflect adult bee diet-driven differences in quality or toxicity of brood food, pupae, or adults upon which *Varroa* feed. Both nutrient limitation and secondary metabolite content have been posited as explanations for poor development of many bee species on Asteraceae pollen [37], as well as for the resistance of Asteraceae specialists to parasitoids [38]. First, the low protein content of sunflower—40% below typical levels for bee-visited plants [36]—could result in brood of small size or poor nutritional quality for *Varroa*. For example, larvae of sunflower pollen-fed bumble bees were 80% smaller than those of *Brassica-*fed bees [61]. Second, *Varroa* may be sensitive to the secondary metabolites in sunflower pollen or nectar [22]. Mites could be directly exposed to secondary metabolites in brood food [56] or by feeding on pupae or adults that have sequestered such metabolites in fat body [62]. Sequestration of secondary metabolites occurs in other insects (e.g., monarch butterflies) and can enhance resistance to parasites [63], predators [64], and parasitoids [65]. The detection of caffeine in *Varroa* excrement after feeding on pupae [66] provides empirical evidence for exposure of mites to these compounds. Third, pollen of sunflower and other Asteraceae is distinguished by an unusual sterol content, including an abundance of Δ-7 sterols that are toxic to some herbivores and a relative lack of 24-methylenecholesterol [67], the main sterol found in honey bees and *Varroa* [68]. Honey bees obtain sterols from pollen, with somatic sterol composition reflecting that of the diet [69]. Incorporation of sunflower sterols into the tissues of pupae or adults could therefore affect *Varroa* survival or reproduction. Further elucidation of physiological and behavioral explanations for sunflower’s effects would facilitate a better understanding of the costs and benefits of sunflower-containing diets, possibly identifying specific components that interfere with mites without harming bees.

Our analysis of planted sunflower acreage in the USA suggests that bees have experienced a progressive decrease in cultivated sunflower availability over the last 40 years (Fig. 3). There are two points of downturn evident in the graph—from 1985-1990 and beginning in the year 2000. Both can be explained by market- and policy-driven shifts in agricultural production in North and South Dakota, which combined for 79% of total acres planted in 2020 and 75% in 2021 (NASS records). The first period of decline likely reflects the implementation of the Conservation Reserve Program, which encouraged conversion of marginal-quality farmland to grassland [71]. This shift was probably beneficial for both honey bees and wild bees, which thrive in CRP lands [12,33]. Because these areas can include considerable amounts of Asteraceae—including wild sunflower [48]—reductions in sunflower cultivation did not necessarily reduce sunflower availability for bees during this period.

**Figure 3.**
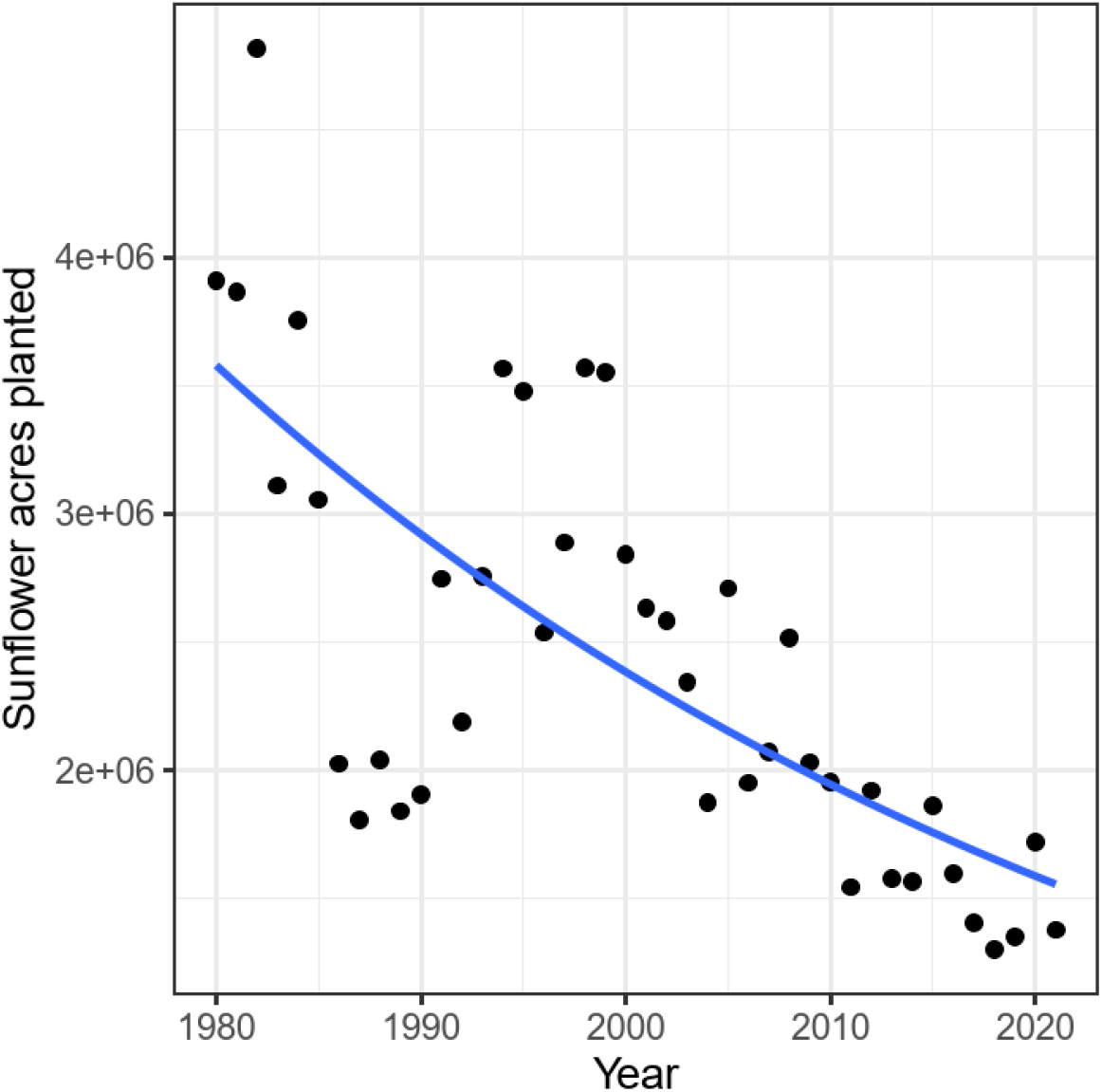
Reductions in sunflower crop area planted in the USA (1980-2021). Points show annual records from the National Agricultural Statistics Service crop production database. Trendline shows Poisson linear model fit. Sunflower crop area has declined by 2.0% per year over the last 4 decades (P < 0.001).

Since 2000, however, there has been a steep decline in both sunflower crop area and Conservation Reserve Program land in the Northern Great Plains [9]. This change has been driven by reductions in CRP acreage quotas and economic incentives for production of biofuel and feed crops [8,45], resulting in displacement of bee-friendly crops and conservation land by corn and soybeans. In North Dakota, sunflower acreage decreased by 35% and CRP-enrolled land area decreased by 27% between 2002 and 2010, concurrent with >50% increases in corn and soy plantings [45]. Between 2006 and 2016, 53% of CRP land surrounding existing North and South Dakota apiaries was converted to crop production, but only 8% were used for bee-friendly crops—as opposed to 67% for corn and soybeans [9]. Because this region is home to not only three-fourths of US sunflowers, but some 40% of US honey bee colonies during the summer as well [8], the reduced availability of sunflowers in this region effectively portrays honey bee access to sunflower in the country as a whole.

The loss of foraging opportunities for bees on wild and cultivated sunflowers could exacerbate susceptibility to mite-induced overwinter collapse that currently plagues beekeepers in the US and worldwide. As a native American species, sunflowers benefit a diverse assemblage of native bees as well [29,30,72], with nectar and pollen of sunflower and its relatives capable of enhancing resistance to other parasites in both social and solitary bees [38–40,42]. Policy-mediated encouragement of sunflower agriculture could therefore provide a win-win solution for managed and wild bees alike, provided that this additional cultivation does not come at the expense of bee-friendly, florally diverse, Asteraceae-rich conservation land [12,33]. Because sunflower yields are enhanced by bee pollination [34] and production of sunflower seed depends on it almost entirely [73,74], sunflower cultivation may provide incentives for pollinator-friendly agricultural and land use practices among growers as well [8]. Whether similar *Varroa* resistance-enhancing benefits result from consumption of wild Asteraceae pollen—for example, *Solidago* spp., whose pollen featured prominently in late-summer pollen collected by honey bees in grasslands [75]—requires further investigation, but seems plausible given the similar antiparasitic effects of sunflower and *Solidago* pollen against *Crithidia bombi* in bumble bees [39].

## CONCLUSIONS

Our results indicate that sunflower enhances honey bee resistance to *Varroa* mites— ectoparasites that drive honey bee colony losses and vector viruses with spillover potential. The timing of sunflower pollen production—which peaks in late summer (in temperate regions), just as mite levels begin to rise towards their peak in October and November [46]—is ideal for reducing levels of infestation during the critical late-season time frame. The current outlook for land use-mediated changes in the status of honey bees and other pollinators has appeared bleak. By contrast, our results illuminate a potential agriculturally compatible intervention to improve honey bee health, while illustrating how policy decisions can shape the availability of antiparasitic resources for pollinators.

## ACKNOWLEDGMENTS

We thank the Dennis vanEngelsdorp and the Bee Informed Partnership for conducting the Maryland field study and virus assessments and coordinating the Honey Bee Disease Survey, George O’Neil for providing colonies for the Massachusetts field experiment, and anonymous reviewers for their service in improving the manuscript.

## DATA AVAILABILITY

All data are supplied in the Supplementary Information, Data S1.

## AUTHORS’ CONTRIBUTIONS

LSA, REI, and JDE conceived the study. LSA, REI, JDE, and BJ designed experiments. BJ, HW, KG, DL, and JDE conducted experiments. YZ, KB, BJ, RM, JVW, JDE, and ECPY analyzed data. ECPY, YZ, BJ, and RM drafted the manuscript. All authors revised the manuscript and gave approval for publication.

## MEDIA PROMOTION

Pollination services of honey bees are threatened by *Varroa* mites and associated pathogens, but current pesticide-based treatments have off-target effects on bees. Enrichment of landscapes with plants that provide medicinal nectar and pollen could provide a more sustainable means of parasite control. We found that sunflower cropland is associated with reductions in mite infestation intensity at the landscape level, and that supplementation of colonies with sunflower pollen reduced infestation by 2.75-fold relative to artificial pollen. Our findings suggest the potential for a simple, agricultural land use-mediated intervention to counteract a main driver of honey bee losses worldwide.

