## Supplementary information for "Mites alight! Sunflower crop area and pollen supplementation enhance honey bee resistance to *Varroa destructor*"

Running title: Sunflower enhances mite resistance in honey bees

Evan C Palmer-Young [1]*, Rosemary Malfi [2], Yujun Zhou [3], Bryanna Joyce [2], Hannah Whitehead [2], Jennifer Van Wyk [2], Kathy Baylis [3], Kyle Grubbs [1], Dawn Lopez [1], Jay D Evans [1], Rebecca E Irwin [4], Lynn S Adler [2]

[1] USDA-ARS Bee Research Lab, Beltsville, MD, USA

[2] Department of Biology, University of Massachusetts Amherst, Amherst, MA, USA

[3] Department of Agricultural & Consumer Economics, University of Illinois at Urbana-Champaign, Urbana and Champaign, Illinois, USA

[4] Department of Applied Ecology, North Carolina State University, Raleigh, NC, USA

.

### EXTENDED METHODS

#### Effects of late-summer sunflower pollen supplementation on field colonies in Maryland

**Pollen treatments.** Honey bee-collected pollens were obtained from commercial suppliers of sunflower (Huading Wax Industries, Henan, China, imported under USDA-APHIS permit number P526P-18-03469) and wildflower (CC Pollen Co., Phoenix, AZ) pollens. Artificial pollen was obtained from a beekeeping supply company (BeePro, Mann Lake Beekeeping, Wilkes-Barre, PA USA). Prior to use in field experiments, bee pollens were sterilized with ethylene oxide at Steris (Isomedix Operations, Inc., Spartanburg, SC) according to a standard protocol (Supplementary Table 1). To ensure that the pollen was fully sterilized, multiple replicates of a bioindicator (*Bacillus atrophaeus*) were embedded into the pollen. Following the sterilization process, the bioindicator replicates were removed from the pollen; no growth of the bioindicator was observed after 7 days' incubation in TSB media at 30-35 °C. Pollens were mixed with sugar water (95 mL 30% sucrose per 450 g pollen) to create a dough-like consistency, divided into 900 g patties, wrapped in wax paper, sealed in plastic, and stored at -20 °C until use.

**Colony maintenance and selection.** Colonies owned and managed by the University of Maryland's Bee Informed Partnership were moved to a common apiary 8 d prior to the start of the study and treated for *Varroa* mites using formic acid strips. Thirty queen-right colonies, containing at least 8 frames of bees each and free of overt disease symptoms (e.g., chalkbrood, foulbrood, wing deformities) were randomly assigned to treatments. One 900 g pollen patty was placed on the topmost brood frame of each colony at weekly intervals for 5 weeks. Each week, the remaining patty from the previous week was collected and weighed to estimate consumption. At baseline and after 5 weeks' treatment, bees (100 mL (~300 individuals) for *Varroa* and *Nosema,* 50 mL for viruses) were collected from a brood frame of each colony.

***Varroa* infestation.** *Varroa* mite levels were determined using standard methods involving 30 min agitation in soapy water followed and straining the resulting water through a 75 mm sieve to collect the mites [1]. Counts were normalized to the number of mites per 100 bees in each sample.

***Nosema* infection.** *Nosema* infection was determined by microscopic spore counts. A 100-bee subsample was crushed with a rolling pin inside a zippered plastic bag (to release gut contents), mixed with 100 mL water, and shaken to homogenize. The number of spores per bee was estimated by hemocytometer counts of a 0.02 µL subsample from a 10 µL aliquot of the mixture.

**Virus infection.** Intensity of infection with seven bee viruses (Acute Bee Paralysis Virus (ABPV), Chronic Bee Paralysis Virus (CBPV), Deformed Wing Virus (DWV), Israeli Acute Paralysis Virus (IAPV), Kashmir Bee Virus (KBV), Lake Sinai Virus-2 (LSBV-2), and Varroa destructor Virus-1 (VDV-1) was determined by quantitative RT-PCR [2].

#### Effects of springtime sunflower pollen supplementation on field colonies in Massachusetts

**Pollen sources.** Pollen was honey bee-collected in August 2017 from two organic sunflower farms (Raine's Honey farm and Matt Messa's Farm) located in Wisconsin, USA. Pollen was collected into bags from pollen traps placed on bee colonies. Trapped pollen was sorted and pooled by batch color into one of three morphotypes: sunflower (approximately 88% sunflower and 12% wildflower), wildflower (approximately 11% sunflower and 89% wildflower) or mix (approximately 28% sunflower and 72% wildflower, based on proportional masses of a 110 g subsample hand-sorted by appearance). Composition of the pollen was confirmed by microscopic examination of fuschin-dyed samples at 400X magnification, which revealed grains with spiked morphology typical of Asteraceae pollen in the bright yellow-orange granules of corbicular pollen. All pollens were sterilized by ethylene oxide (Supplementary Table 1). Pollen patties (170 g each) were prepared by homogenization of pollen in a blender, followed by mixing with sucrose solution (95 mL 30% sucrose per 450 g pollen) to create a dough-like consistency. All patties were wrapped in wax paper, sealed, and stored at -20 °C until use.

**Colony selection.** We selected 55 colonies (11 per treatment) originally located at an apiary in Barre, Massachusetts, USA (Autumn Morning Rain Farm (42.439486, -73.989308)). Colonies were randomly assigned to treatments prior to relocation to one of five sites (n = 2-3 colonies per treatment and site). The five sites were: Barre Rd orchard: (42.3312, -72.1145), McEvoy Rd. Farm: (42.3523, -72.1293), Petersham Rd. Farm: (42.3936, -72.2027), Cranberry Bog: (42.3418, -72.158) and the original Autumn Morning Rain Farm location. Each site was separated by >3.2 km to ensure spatial independence. We selected colonies with at least four full frames of bees, excluding those missing a queen or exhibiting abnormal queen behavior. Each week, a fresh pollen patty was placed on the topmost brood-containing box to encourage consumption. Consumption of the previous week's patty was estimated visually to the nearest 5%. One colony that exhibited consistently poor consumption (<25% consumption in each of the first three weeks) was excluded from statistical analyses *a priori*.

**Colony observations and sample collection.** Before starting treatments, colonies were visually assessed to confirmed that all were queenright and free from brood diseases. Colonies were sized by counting the number of hive frames occupied by bees [3]. These assessments were repeated after two and five weeks' treatment. *Varroa* and *Nosema* were measured pre- and post-treatment (0 and 5 weeks). Viruses were assessed post-treatment only.

**Parasite quantification.** To assess the level of mite infestation, we sampled 120 mL of adult bees (approximately 300) from each colony, using equal quantities from an inner (brood-filled) and outer frame. Samples were agitated (2 min) in 95% ethanol; mites were counted in the resulting liquid. These samples were then stored in ethanol at 4 °C until homogenization for *Nosema* analysis by microscopy (see *Nosema* infection above). For post-treatment viral analysis, live worker bees (~250 individuals, collected from brood frames after six weeks' treatment) were shipped overnight to the Bee Informed Partnership at the University of Maryland in boxes (Mann Lake Riteway Shippers) outfitted with a 90 mL petri dish filled with "queen candy" (6:1 powdered sugar: corn syrup blend) and a 15 mL water tube for sustenance. Seven viruses were quantified by qPCR using the same methods employed for the National Honey Bee Disease Survey [4]. The "mix" treatment was excluded from viral analysis, as it had no counterpart in the late-summer supplementation experiment.

#### Effects of sunflower pollen on caged bees

Mortality was minimal during the trial; surviving bees were stored at -80$^{o}$ C until RNA extraction.

**Quantification of infection.** RNA was extracted from a pooled sample of 20 bees per cup using TRIzol reagent (Thermo Fisher Scientific), as previously described in [5]. Briefly, bees were ground in sterile plastic bags using an RNAse-inhibiting lysis buffer and pelleted in a microcentrifuge at 5000 rpm for 1 min. The aqueous solution was transferred to a new sterile tube, to which 500 µl TRIzol was added. After mixing, 200 µl chloroform was added to each tube, products were again mixed and the aqueous phase was added to 500 µl isopropanol. After mixing and incubation at room temperature for 10 min, RNA was pelleted by centrifugation (10 min, 12,500 rpm) at 4°C. The supernatant was removed and the pellets washed with 1 ml of cold 75-80% nuclease-free ethanol, re-spun at 4°C for 5 min at full speed, and all ethanol removed. Pellets were dissolved in 100 µl of RNase-free water.

RNA extracts were treated with DNase I at 37 °C for 1 h followed by 10 min at 75 °C. First-strand complementary DNA (cDNA) was generated from 2 µg total RNA using a master mix containing 50 U Superscript II (Invitrogen), random primer set (7-mer at 10 mM concentration), 2 nmol dNTP mix, 2 nmol polydT-18, and 0.1 nmol polydT (12–18). The cDNA synthesis was carried out at 42 °C for 50 min followed by 15 minutes at 70 °C. Pathogen loads were estimated via quantitative real-time PCR (1 ul cDNA template in a 20 µlreaction) using Bio-Rad SsoFast™ SYBR® Green Supermix, 96-well optical PCR plates, and a Bio-Rad CFX Connect™ thermal cycler. DWV was assessed using primers DWV.F (GAGATTGAAGCGCATGAACA) and DWV.R (TGAATTCAGTGTCGCCCATA); VDV-1 using primers VDV.F (GCCCTGTTCAAGAACATG) and VDV.R (CTTTTCTAATTCAACTTCACC); *Nosema ceranae* using Ncer.F (CGTTAAAGTGTAGATAAGATGTT) and Ncer.R (GACTTAGTAGCCGTCTCTC); and trypanosomatids using primers Trypan1.F (GTGCAGTTCCGGAGTCTTGT) and Trypan1.R (CTGAGCTCGCCTTAGGACAC). mRNA's for honey bee actin (gene GB44311) and Ribosomal protein S5 (RpS5) were used to normalize infection intensity, with primers Actin.F (TTGTATGCCAACACTGTCCTTT) and Actin.R (TGGCGCGATGATCTTAATTT) and RpS5.F (AATTATTTGGTCGCTGGAATTG) and RpS5.R (TAACGTCCAGCAGAATGTGGTA) [6].

Quantitative real-time PCR was carried out in duplicate on a Bio-Rad (Hercules, CA, USA) CFX-96 thermal cycler using a profile of 95 °C for 30 seconds, followed by 50 cycles of denaturation at 95 °C for 5 seconds and annealing/extension 60 °C for 30 seconds. Positive controls were included on each plate for each target, along with no-template negative controls.

**Supplementary table 1. Procedure for pollen sterilization by ethylene oxide (EO).**

| **Cycle phase** | **Setpoint** |
|---|---|
| Chamber temperature | |
| a. Cycle start temperature | 39 °C |
| b. Exposure dwell temperature | 39 °C |
| Initial evacuation A | |
| a. Evacuation pressure | 1.0 inHgA |
| b. Evacuation rate | 1.0 inHgA/min |
| Nitrogen dilution (pre-EO exposure) | |
| a. Injection pressure | 15.0 inHgA |
| b. Injection rate | 1.0 inHgA/min |
| c. Evacuation pressure | 14.9 inHgA |
| d. Evacuation rate | 1.0 inHgA/min |
| e. Vacuum hold time | 60 min |
| f. Number of repetitions | 1 |
| Initial evacuation B | |
| a. Evacuation pressure | 1.0 inHgA |
| b. Evacuation rate | 1.0 inHgA/min |
| Humidity infection B | |
| a. Injection pressure | 1.7 inHgA |
| Humidity dwell | |
| a. Humidity dwell control pressure | 1.7 inHgA |
| b. Dwell time | 60 min |
| Sterilant injection | |

| | |
|---|---|
| a. Vaporization water temperature | 75 °C |
| b. Infection pressure | 13.4 inHgA |
| c. Injection rate | 1.0 inHgA/min |
| Sterilant exposure | |
| a. Sterilant dwell control pressure | 13.4 inHgA |
| b. Injection rate | 1.0 inHgA/min |
| c. Dwell time | 240 min |
| Sterilant removal | |
| a. Evacuation pressure | 1.0 inHgA |
| b. Evacuation rate | 1.0 inHgA/min |
| Nitrogen Wash (post-EO exposure) | |
| a. Injection pressure | 20 inHgA |
| b. Injection rate | 1.0 inHgA/min |
| c. Evacuation pressure | 1.0 inHgA |
| d. Evacuation rate | 1.0 inHgA/min |
| e. Number of repetitions | 8 |

### SUPPLEMENTARY DATA

Raw data are provided in Supplementary Data S1 for the landscape study (“landscape_dataset”), late-summer supplementation experiment in Maryland (‘Maryland_supplementation”), springtime supplementation experiment in Massachusetts (“Massachusetts_supplementation”), caged bee experiment (“Cage_trials”), and US sunflower crop area records (“Sunflower_US_totals”). Each csv file is accompanied by a text file with variable definitions.
