## Supplementary material for "Mites alight! Sunflower crop area and pollen supplementation enhance honey bee resistance to *Varroa destructor*": Cage_trials_metadata.txt

#Metadata:
#Sample: unique sample ID
#Pollen.sunflower.pct: proportion sunflower in pollen treatment
#Target: parasite quantified
#Ct: mean cycle time of two technical replicates
#Actin, RPS5: cycle times of the two control genes
#Mean.controls: mean of Actin and RPS5
#dCt: normalized infection intensity, 
 #i.e. difference in cycle times of control genes (Mean.controls) and target (Ct)
